# Resistance to the antifungal activity of Aprotinin occurs through mutations in genes that function in cation homeostasis

**DOI:** 10.1101/2020.06.22.164863

**Authors:** Amanda I. McColl, Rohan G.T. Lowe, James A. McKenna, Marilyn A. Anderson, Mark R. Bleackley

**Author notes:** NCBI SRA accession number: *S. cerevisiae* (R64-1-1.23) reference genome: PRJNA43747, NaD1-R and Controls: PRJNA434021, BPTI-R and Controls: SUB7621764. Corresponding author: Marilyn A. Anderson, (03) 9479 1255.

## Abstract

An increase in the prevalence of fungal infections is coinciding with an increase of resistance to current clinical antifungals, placing pressure on the discovery of new antifungal candidates. One option is to investigate drugs that have been approved for use for other medical conditions that have secondary antifungal activity. Aprotinin, also known as Bovine Pancreatic Trypsin inhibitor (BPTI), is an antifibrinolytic that has been approved for systemic use in patients in some countries. Bleackley and coworkers (2014) revealed that BPTI also has antifungal activity against *S. cerevisiae* and *C. albicans* and does this by targeting the magnesium transporter *ALR1.* Here we have further investigated the potential for aprotinin to be used as an antifungal by assessing the development of resistance. We used an *in vitro* model to assess the evolution of BPTI resistance/tolerance whereby BPTI was serial passaged with the model organism *S. cerevisiae.* Resistance to BPTI developed more quickly than resistance to the plant defensin NaD1 and the clinical antifungal, caspofungin. Full genome sequencing of resistant lines revealed that resistance to BPTI developed as the result of a deleterious mutation in either the *ptk2* or *sky1* genes. This revealed that cation homeostasis and transport functions were particularly affected in *S. cerevisiae* after exposure to BPTI. Therefore, the mutations in these genes probably decreases release of magnesium and other cations from the cell, protecting the yeast from the limiting intracellular magnesium levels that arise when BPTI blocks the magnesium transporter Alr1p.

## Introduction

Aprotinin, also known as bovine pancreatic trypsin inhibitor (BPTI), has been used as a drug to reduce bleeding during cardiopulmonary bypass (CPB) and was sold by Bayer under the trade name Trasylol (Bidstrup *et al*, 1989). Aprotinin slows down fibrinolysis, the process that leads to the breakdown of blood clots (Royston *et al*, 1987). In 1987, a small study was published that reported a reduction of blood loss and need for transfusions in patients treated with Aprotinin which resulted in wide use of the drug in cardiac, hepatic and orthopedic surgery (Royston *et al.*, 1987). However, subsequent large scale clinical studies revealed potential safety concerns associated with the use of Aprotinin, including an increased risk of renal failure, myocardial infarction, heart failure, stroke, and encephalopathy (Karkouti *et al*, 2006; Mangano *et al*, 2006). As a result, marketing of Aprotinin was suspended in 2007, but the results of these studies have been challenged and in 2012 Aprotinin was reintroduced in Canada and the European Medical Association also recommended a lift of the ban (Furnary *et al*, 2007; McMullan & Alston, 2013). Therefore, it may be time to rethink the potential uses for Aprotinin, including those outside of reducing blood loss during surgery.

In a previous study, Bleackley and co workers, discovered that Aprotinin (BPTI), also inhibits the growth of *Saccharomyces cerevisiae* and the human fungal pathogen *Candida albicans* (Bleackley *et al*, 2014). It does this by binding to the magnesium transporter Alr1p, blocking magnesium uptake into the cell and causing cell cycle arrest (Bleackley *et al.*, 2014). It also works synergistically with the well characterised antifungal plant defensin, NaD1 against *Fusarium graminearum*, *Colletotrichum graminicola* and *Candida albicans*, although the exact mechanism of synergy is unclear (Bleackley *et al*, 2017). Aproptinin can be administered to patients via injection under strict guidelines and benefit-risk assesment, therefore aprotinin is a potential treatment for systemic fungal infections that requires further investigation. One important consideration in the use of a drug to treat microbial infections is the potential for the pathogenic microbes to develop increased tolerance and eventually, resistance to the drug.

In this study, we generated yeast strains that have increased tolerance to Aprotinin (BPTI), and identified the genetic mutations linked to the decreased sensitivity to BPTI. We found that resistance to BPTI developed more quickly than resistance to the plant defensin NaD1 and to the clinical antifungal caspofungin. Resistance was caused by single point mutations that inactivated either the *sky1* or *ptk2 genes* that function in ion transport. These mutations are likely to increase magnesium accumulation in the cells and prevent cell cycle arrest. We did not find mutations in the gene encoding the magnesium transporter Alr1p that BPTI is known to bind to. Interestingly synergy between NaD1 and BPTI was not affected in yeast strains that were more resistant to either one of the two peptides. That is, sensitivity to each peptide was restored by the addition of subinhibitory levels of the synergy partner.

## Material and methods

### Fungal Strains

The *S. cerevisiae* strain BY4741 *(MATa his3Δ0 leu2Δ0 met15Δ0 ura3Δ0)* was purchased from Thermo Scientific. Single deletion strains were retrieved from the haploid non-essential deletion collection (Thermo Scientific)(Winzeler *et al*, 1999). Production of NaD1 and caspofungin-resistant lines of *S. cerevisiae* are described in (McColl *et al*, 2018).

The *C. albicans* strain ATCC 90028 was purchased from In Vitro Technologies Pty. Ltd. *C. neoformans* KN99 was purchased from The Fungal Genetics Stock Centre. The yeasts *S. cerevisiae, C. albicans and C. neoformans* were routinely cultured on YPD medium at 30°C *A. fumigatus* ATCC MYA-3626 was purchased from In Vitro Technologies Pty. Ltd.

A patient isolate of *T. rubrum* was acquired from the National Mycology Reference Centre (Adelaide, SA). The filamentous fungi *A. fumigatus* and *T. rubrum* were cultured at 28°C on V8 agar.

### Antifungal molecules

Aprotinin (BPTI) was purchased from Astral Scientific (Australia), NaD1 and NaD2 were purified from *Nicotiana alata* flowers as described in ((Dracatos *et al*, 2014; Lay *et al*, 2003). Caspofungin was purchased from Sigma (Australia).

LL-37 was synthesized by GL Biochem (China) and Bac2a was synthesized by GenScript (China).

### Culturing in the presence of antifungal molecules to develop resistance

The culturing method to develop resistance to BPTI was performed as described in (McColl *et al.*, 2018) whereby, *S. cerevisiae* BY4741 was grown overnight at 30°C with agitation in 5 mL of yeast extract-peptone-dextrose (YPD). The overnight culture was then diluted to an OD600 nm of 0.01 in 50% strength potato dextrose broth (½ PDB) before addition of antifungal molecules. Cultures were initially grown with the BPTI at 0.5x the mimimum inhibitory concentration (MIC) (5 μM) or 1x MIC (10 μM) alongside a negative control lacking antifungals. Three independent lines for the BPTI resistant and untreated controls were grown in parallel. The cultures were incubated overnight at 30°C with agitation. The cultures that exhibited growth at the highest concentration of BPTI were then used to seed cultures at a higher concentration of BPTI. Sub-culturing was stopped once growth occurred at 32 times the original MIC.

### Single Colony Isolation of Resistant Strains

Cultures resistant to the antifungal molecule were streaked out to isolate single colonies on non-selective YPD agar. Three colonies were picked from each line and their resistance was re-tested, the colony with the highest resistance to the antifungal was used in subsequent experiments.

### Antifungal Assays

Antifungal assays were performed as described in (Hayes *et al*, 2013). Briefly, yeast cultures *S. cerevisiae, C. albicans and C, neoformans* were grown overnight (30°C, 250 rpm) in 5 mL YPD and diluted to an OD600 of 0.01 in ½ PDB. Antifungal molecules were prepared at 10x the assay concentration and 10μL was mixed with 90 μL of diluted yeast culture before incubation for 24h at 30°C.

Filamentous fungi *T. rubrum and A. fumigatus* were cultured for 3 weeks (28°C) on V8 agar. To prepare the inoculum for the filamentous fungi, the spores were obtained by flooding plates with sterile water. Hyphal matter was removed by filtration through sterile facial tissue. Spores were quantified by hematocytometer and diluted to 50,000 spores/mL in 1/2 PDB before use. Antifungal molecules were prepared at 10x the assay concentration and 10μL was mixed with 90 μL of diluted fungal spores before incubation for 48 h at 28°C.

The final OD600 was measured using a SpectraMAX M5e plate reader (Molecular Devices).

### Synergy Assays

Synergy assays were performed as described in (Bleackley *et al.*, 2017) whereby, *S. cerevisiae* was cultured in liquid YPD overnight at 30°C beforecells were diluted to an OD600 of 0.01 in ½PDB. Antifungals were prepared at 10 times the assay concentration.

Aliquots of the test molecules (10 μl) were placed into the wells of a 96-well microtiter plate in a standard checkerboard array. Cells (80 μl) were then added to all the wells of the plate followed by incubation at 30°C for 24 h. Growth was monitored by measuring the optical density at 595 nm using a SpectraMax M5e plate reader (Molecular Devices). Synergistic interactions were identified using the fractional inhibitory concentration (FIC) calculation with a cutoff of 0.5 indicative of synergy (FIC = MIC_A_combination/MIC_A_alone + MIC_B_combination/MIC_B_alone where MICAcombination is the MIC of agent A in combination and MICAalone is the MIC of agent A alone) as described previously by (Bleackley *et al.*, 2017). If a test molecule was not inhibitory when used in isolation, the MIC was set to two times the top concentration tested.

### Cell Growth Assay

*S. cerevisiae* BY4741 cultures were grown overnight (30°C, 250 rpm) in 5 mL of YPD and were diluted to an OD_600_ of 0.5 in 1 mL YPD or ½ PDB before 100 μL was dispensed into each of 8 wells of a 96 well microtiter plate. The plate was then incubated in a SpectraMAX M5e plate reader (Molecular Devices) at 30 °C. Optical density at 600nm was recorded for each well every 30 min over the 48h culture period and averaged across the 8 replicates.

### Stress assay with hydrogen peroxide, calcofluor white, NaCl and SDS

Hydrogen peroxide (0.625, 1.25, 2.5 or 5 mM), calcofluor white (1, 2.5, 5 or 10 μg), NaCl (100, 200 or 300 mM), or SDS (12.5, 25, 50 or 100 μg) were diluted in YPD agar media (25 mL) just before the plates were poured. Yeast cultures were grown overnight in 5mL of YPD before they were diluted to an OD600nm of 0.1 in 1 mL of MilliQ-purified water. A five-fold dilution series of each yeast strain was spotted (4 μL per spot) onto the agar plates with the added stress factors and incubated overnight at 30°C. Plates were imaged using a ChemiDoc (Biorad).

### Stress Assay with Ultraviolet Light

*S. cerevisiae* cultures were grown overnight in 5 mL YPD and diluted to an OD600nm of 0.1 in 1 mL MilliQ-purified water. A five-fold dilution series of each strain (4 μL per spot) was then spotted onto a YPD agar plate and allowed to dry before exposure to UV light (Phillips, 30w bulb at 50 cm) for 1.2, 2.4, 5.2 or 10.4 min.

### DNA extraction from wild-type and resistant strains of *S. cerevisiae*

Genomic DNA was extracted using the Qiagen DNeasy^®^ plant mini kit. Three independent lines of BPTI-resistant strains and three lines of the no-treatment controls were sequenced. Sequencing was completed at the La Trobe Genomics Platform, using the Illumina MiSeq V3 chemistry. One run was performed for all 6 strains, generating 25 million 300 bp paired-end reads. The pre-processing and variant discovery steps were performed as described by the GATK best practices as summarized in (McKenna *et al*, 2010).

### Genomic analysis of resistant strains of *S. cerevisiae*

Genomic analysis was completed as described in (McColl *et al.*, 2018) and was as follows:

### Sequence pre-processing

The raw sequence reads were converted to Sam format and illumine adapters were identified using Picard tools (v.2.4.1) fastqtosam and markilluminaadapters. The reads were then aligned to the *S. cerevisiae* (R64-1-1.23) reference genome using BWA-mem (v0.7.12) (Engel *et al*, 2014). The aligned read files were merged using Picard (v.2.4.1) mergebamallignment and subjected to quality control and filtering using Picard (v2.4.1) markduplicates and GATK (v.3.6) realignertargetcreater and baserecalibrator.

### Variant Discovery

Variations that were either SNVs (single-nucleotide variants) or INDELs (insertion/deletion) were discovered using GATK (v.3.6) Haplotypecaller (Sherry *et al*, 2001). The samples were merged using GATK (v.3.6) combinegvcf and then joint genotyping was performed using GATK (v.3.6) GenotypeGVCFs. SNV’s and indels were extracted based on default quality parameters using GATK (v.3.6) VariantFiltration and VariantRecalibrator.

### Variant Refinement

Variants were annotated using SnpEff (v.2.4) (Cingolani *et al*, 2012). VEP (variant effect predictor) marked any codon changes as either tolerant or deleterious (McLaren *et al*, 2016). SnpSift (v.2.4) was used to identify SNVs or indels that were present in the BPTI-resistant replicates and not in the Control strains. The variants were then inspected manually using IGV (v.2.3.77) (Robinson *et al*, 2011).

### Data Availability statement

Strains are available upon request. The authors affirm that all data necessary for confirming the conclusions of this article are represented fully within the article and its tables and figures. The publically available *S. cerevisiae* (R64-1-1.23) reference genome was obtained from the NCBI SRA database with accession number PRJNA43747. Raw sequencing data for NaD1-R and Controls from (McColl *et al.*, 2018) are publicly available from the NCBI SRA database with accession number PRJNA434021. BPTI-R sequencing data has been uploaded to the NCBI SRA database with the submission number SUB7621764 and is confidentially available to the editors and reviewers upon request. This submission will be made publically available once paper is accepted.

## Results

### BPTI inhibits growth of *S. cerevisiae, C. albicans and C. neoformans*

The antifungal activity of BPTI was tested in a growth assay with the yeast *S. cerevisiae, C. albicans* and *C. neoformans* as well as the filamentous fungi *T. rubrum* and *A. fumigatus.* BPTI was effective against *S. cerevisiae* with an MIC of 12.5 μg/mL and *C. neoformans* with an MIC of 9.4 μg/mL (Table 1). It was less effective against *C. albicans* with an MIC of 100 μg/mL and was ineffective against both filamentous fungi, *T. rubrum* and *A. fumigatus* at concentrations up to 200 μg/mL (Table 1).

**Table 1.** BPTI is effective against yeast but not filamentous fungi. The activity of BPTI was tested against *S. cerevisiae, C. neoformans, C. albicans, T. rubrum* and *A. fumigatus.* BPTI was more effective against *S. cerevisiae* and *C. neoformans* than *C. albicans* and was not effective against *T. rubrum* or *A. fumigatus.* Standard error is based on 2 biological replicates for each yeast and 1 biological replicate for filamentous fungi.

| Strain | Rep 1 MIC (µg/mL) | Rep 2 MIC (µg/mL) | Average MIC (µg/mL) |
| --- | --- | --- | --- |
| *S. cerevisiae* | 12.5 | 12.5 | 12.5 |
| *C. neoformans* | 12.5 | 6.25 | 9.4 |
| *C. albicans* | 100 | 100 | 100 |
| *T. rubrum* | >200 | N/A | >200 |
| *A. fumigatus* | >200 | N/A | >200 |

### Evolution of resistance to BPTI

Yeast strains with increased tolerance to BPTI were developed by continuous culture of *S. cerevisiae* at sub-inhibitory concentrations. Each time the MIC increased; the dose of antifungal was doubled. The starting concentration of BPTI in the continuous cultures was 5 μM. After 12 rounds of sub-culturing strains were able to grow in 160 μM BPTI, a 32-fold increase in the MIC (Figure 1A).

**Figure 1.**
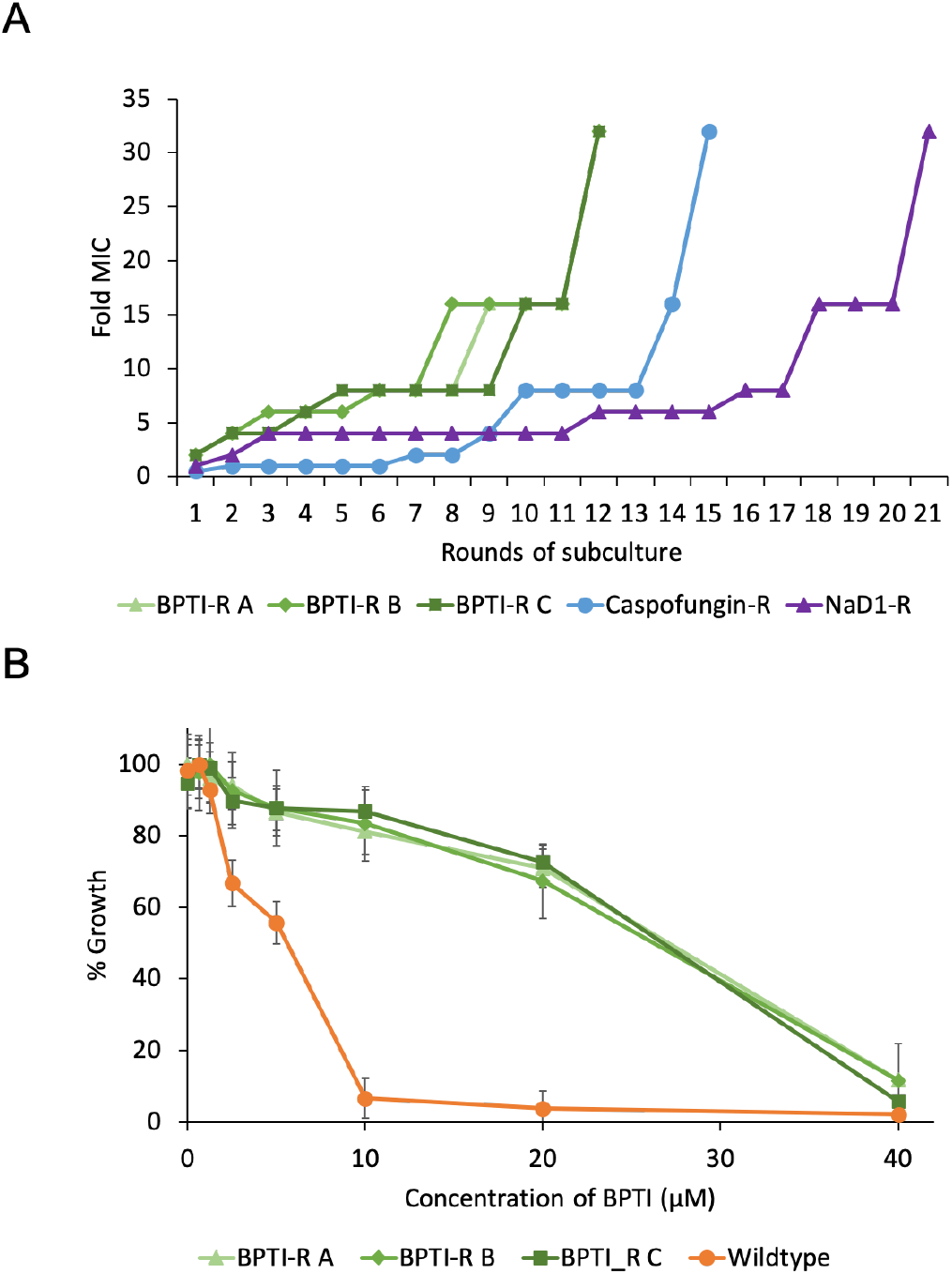
(A) Resistance to BPTI develops more quickly than resistance to caspofungin or NaD1. Three independent cultures of BPTI-resistant yeast are shown, along with a representative example of the caspofungin and NaD1-resistant lines described in our previous paper (McColl *et al.*, 2018). Resistance to BPTI developed steadily with the MIC increasing 32-fold after 12 rounds of growth. It took 15 rounds of subculturing to produce an equivalent level of resistance to caspofungin and 21 rounds for NaD1. **(B) Confirmation of BPTI resistance.** Growth of BPTI-resistant strains of *S. cerevisiae* once selection pressure was removed and independent isolates were selected, at various concentrations of BPTI compared to the BPTI inhibition of the wildtype *S. cerevisiae* control. BPTI-resistant strains were 4-fold more resistant than wildtype. Growth % is relative to the untreated control for each strain. Error bars represent +/− standard error of the mean *(n* = 3).

Once selection pressure was removed, the MIC of the resistant strains can decrease. Therefore, the three BPTI-resistant cultures were streaked onto agar and a pure strain for each culture was isolated. Their resistance phenotype was confirmed using a standard antifungal assay (Figure 1B). These strains were used for all further experimentation. The BPTI-resistant strains were 4-fold more tolerant to BPTI than wildtype, with an MIC of 40 μM compared to the MIC of 10 μM for wildtype (Figure 1B).

### Resistance to BPTI confers resistance to some but not all antifungal peptides

We examined whether the enhanced tolerance of the BPTI resistant strains was broadspectrum or specific to BPTI by comparing the susceptibility of BPTI-resistant lines and wildtype to a set of antimicrobial molecules. The BPTI-resistant strains were more resistant to the human cathelicidin LL-37 with an MIC of 20 μM compared to wildtype inhibition at 5 μM (Figure 2A) and they were also more resistant to NaD2, another plant defensin from *Nicotiana alata,* with an MIC of 20 μM compared to 10 μM for the wild type (Figure 2B). However, this enhanced tolerance did not extend to all AFPs tested. NaD1 (Figure 2C) and caspofungin (Figure 2D) had similar levels of activity against the BPTI resistant strains as the wild-type strain and BPTI-resistant strains were more sensitive than wildtype strains to the peptide Bac2a, which is a loop swap variant of bactenecin, a bovine cathelicidin (Figure 2E).

**Figure 2.**
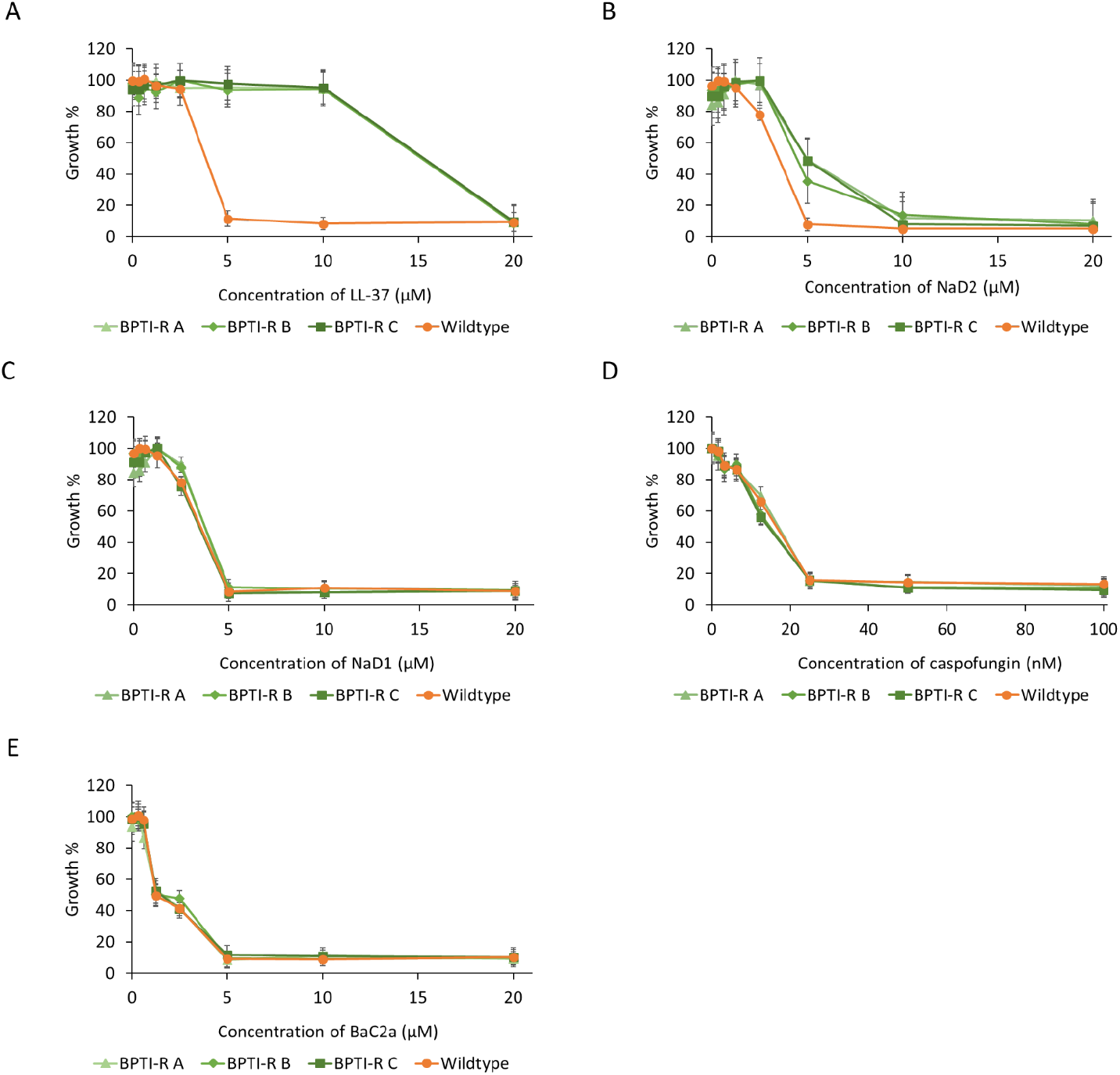
Resistance to BPTI is not broad spectrum. Growth inhibition of BPTI-resistant strains by a selection of antimicrobial peptides of different origin and mechanisms of action. The peptides examined were **(A) LL-37, (B) NaD2, (C) NaD1, (D) caspofungin** and **(E) Bac2a**. BPTI-resistant strains were more tolerant than the wildtype cells to the antifungal peptides LL-37 (A) and NaD2 (B), but not to the antifungals NaD1 (C), caspofungin (D) and Bac2a (E). Growth % is relative to the untreated control for each strain. All experiments were performed in triplicate. Error bars represent +/− one standard error of the mean (*n*= 3).

### BPTI-resistance is associated with cation homeostasis and cell wall stress

As the BPTI-resistant strains displayed differential sensitivity to AMPs, we assessed whether the BPTI-resistant strains differed to wildtype in susceptibility to a selection of abiotic stresses. The BPTI-resistant strains grew better than wildtype at elevated NaCl concentrations (Figure 3A). Similarly, BPTI-resistant strains were more resistant to calcofluor white (CFW) than wildtype yeast. BPTI-resistant strain B was also more tolerant to hydrogen peroxide compared to wildtype cells but not BPTI resistant strains A and C which had the same sensitivity as wildtype cells (Figure 3A). BPTI-resistant strains and wildtype cells were equally sensitive to UV and to SDS (Supplementary figure A).

**Figure 3.**
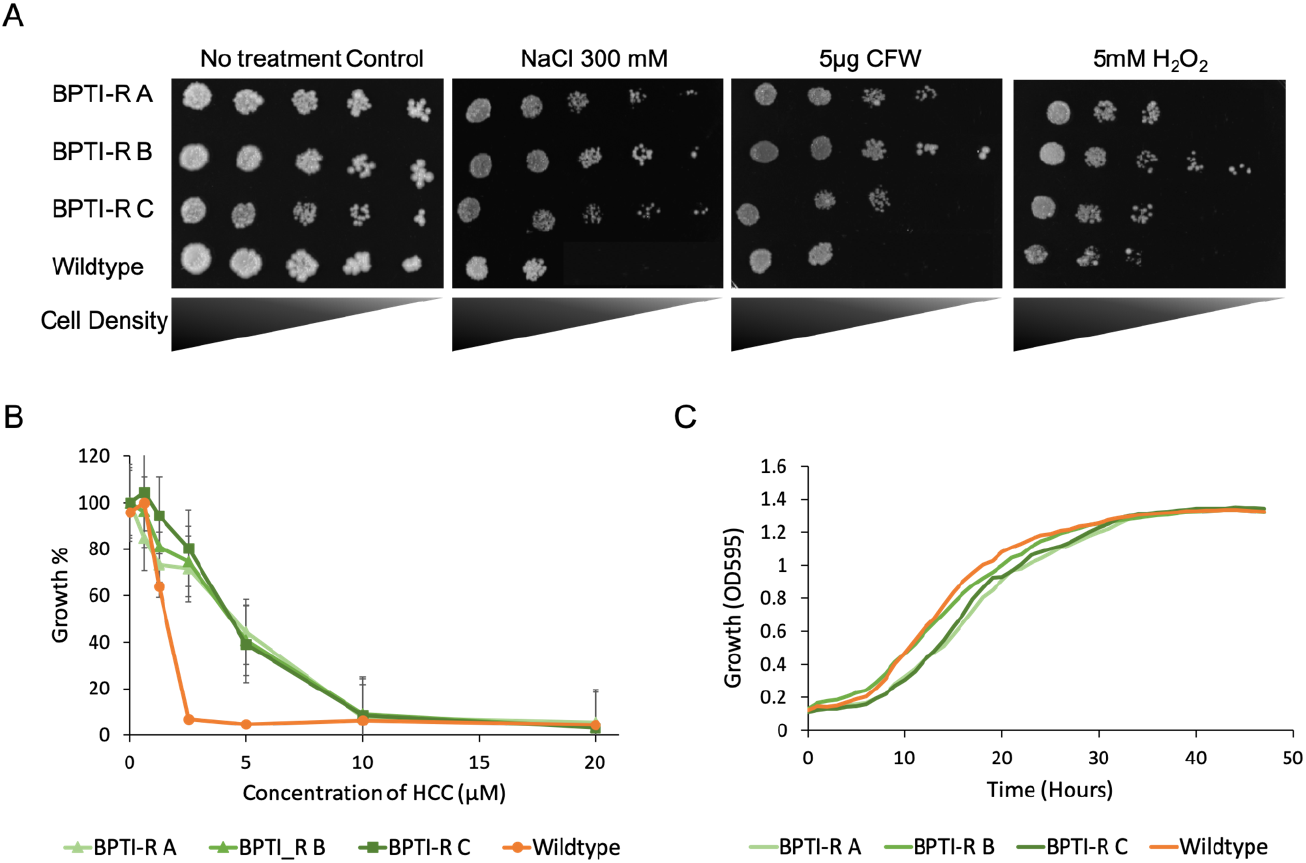
Fitness of BPTI resistant strains in the presence of various abiotic stressors. **(A) BPTI-resistant strains were more tolerant to sodium chloride, calcofluor white and hydrogen peroxide than wildtype cells.** BPTI-resistant and wild-type *S. cerevisiae* BY4741 cells were diluted and spotted out on to YPD Agar containing different concentrations of abiotic stressors. A no treatment control was prepared using the same cell preparations and at the same time as the treatment plates. The three BPTI-resistant strains grew better than wildtype in the presence of 300mM of NaCl BPTI-resistant strains were more resistant to 5μg/mL CFW compared to wild type. BPTI-resistant strain B grew better than wildtype in the presence of 5mM hydrogen peroxide whereas BPTI-resistant strains A and C had similar growth to wildtype. Images are representative of three replicate experiments. **(B) BPTI-resistant strains were more tolerant to Hexamine (III) cobalt chloride than wildtype.** Growth inhibition of BPTI-resistant strains by Hexamine (III) cobalt chloride (HCC) relative to wildtype control. BPTI-resistant strains were more tolerant to the magnesium channel inhibitor HCC with an MIC of 10 μM compared to wild type at 2.5 μM. Growth % is relative to the highest measured absorbance for each strain and the wild-type *S. cerevisiae* BY4741. Error bars represent +/− standard error of the mean *(n=* 3). **(C) Cell growth of BPTI-resistant strains in YPD media compared to wild-type *S. cerevisiae* BY4741.** There was no difference in growth of the BPTI-resistant strains in YPD compared to wildtype.

We reported in an earlier paper that BPTI blocks the magnesium transporter Alr1p and restricts growth of *S. cerevisiae* (Bleackley, 2014). Therefore, we assessed whether the BPTI-resistant strains in this study had defects in magnesium transport by examining whether they were more tolerant to Hexamine (III) cobalt chloride (HCC), a magnesium channel inhibitor. The BPTI-resistant strains were 4-fold more tolerant to HCC with an MIC of 10 μM HCC compared to wildtype which was inhibited at 2.5 μM HCC (Figure 3B)

The fitness of the BPTI-resistant strains was assessed relative to wildtype using media with no antifungal molecules. The growth rates of the BPTI-resistant strains were similar to the wildtype, with strains A and C growing slighter slower (Figure 3C).

### Genetic characterisation of BPTI resistance

Mutations associated with BPTI resistance, were identified by sequencing the genomes of the BPTI-resistant and non-selected control lines. Five genes (Erg3 Ser24Lue, Gda1 Cys462Phe, Nrp1 Asn444Ser, Ptk2 Gly469*stop, Sky1 Gln65*stop) had mutations in their protein coding regions leading to amino acid substitutions or truncated proteins. Four other genes (Gex1, Sok1, tT(XXX)Q2, YBR298C-A) had mutations in the upstream non-coding region (Table 2). A description of the predicted functions of the affected genes is also listed in Table 2.

**Table 2.**
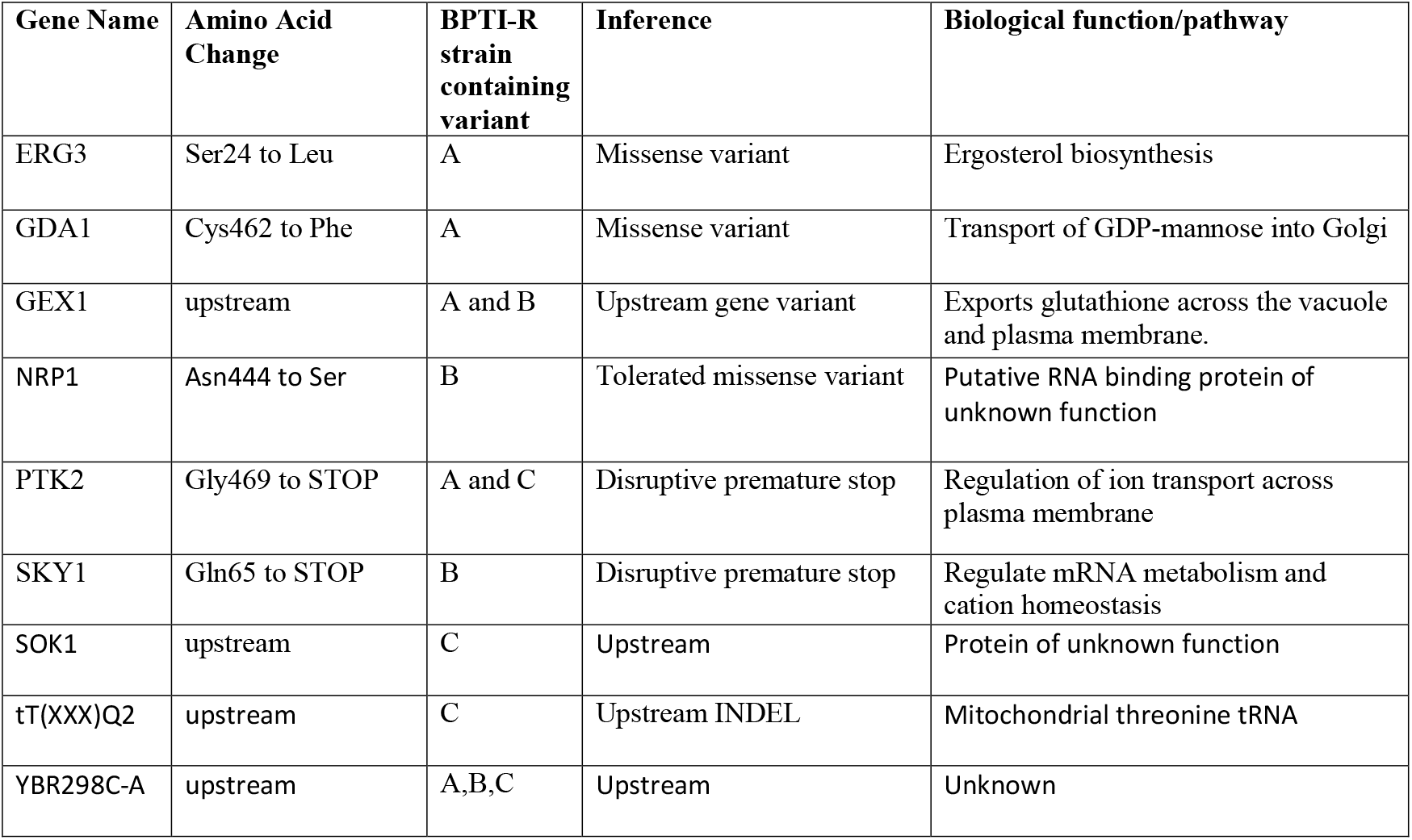
Summary of variants present in BPTI-R strains. The genes that have mutations linked to BPTI resistance and the description of their role in *S. cerevisiae.* The program Variant effect predictor was used to discover the impact of mutations on gene function. Variants were selected if mutations impacted the protein coding region and were present in the resistant strains and absent from the control strains. Upstream gene variants were not selected for further analysis. These genes may be viewed on the Saccharomyces Genome Database www.yeastgenome.org (Cherry *et al*, 2012; Engel *et al.*, 2014).

| Gene Name | Amino Acid Change | BPTI-R strain containing variant | Inference | Biological function/pathway |
| --- | --- | --- | --- | --- |
| ERG3 | Ser24 to Leu | A | Missense variant | Ergosterol biosynthesis |
| GDA1 | Cys462 to Phe | A | Missense variant | Transport of GDP-mannose into Golgi |
| GEX1 | upstream | A and B | Upstream gene variant | Exports glutathione across the vacuole and plasma membrane. |
| NRP1 | Asn444 to Ser | B | Tolerated missense variant | Putative RNA binding protein of unknown function |
| PTK2 | Gly469 to STOP | A and C | Disruptive premature stop | Regulation of ion transport across plasma membrane |
| SKY1 | Gln65 to STOP | B | Disruptive premature stop | Regulate mRNA metabolism and cation homeostasis |
| SOK1 | upstream | C | Upstream | Protein of unknown function |
| tT(XXX)Q2 | upstream | C | Upstream INDEL | Mitochondrial threonine tRNA |
| YBR298C-A | upstream | A,B,C | Upstream | Unknown |

**Table 3.**
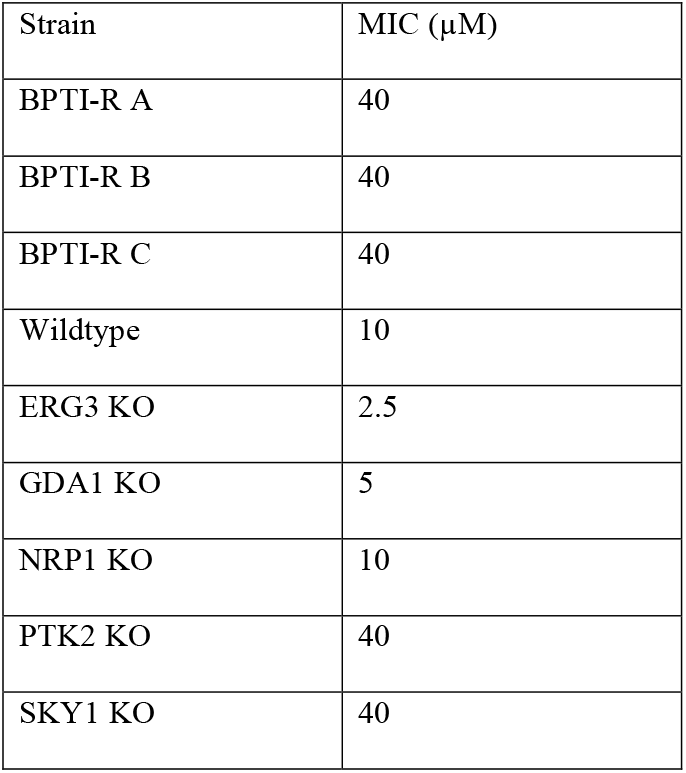
Comparison of BPTI activity against single-gene deletion strains representing key resistance variants. The activity of BPTI against BPTI-resistant strains A, B and C, wild-type *S. cerevisiae* BY4741 and the gene knock outs of ERG3, GDA1, NRP1, PTK2 and SKY1 in ½ PDB. This is a representative example of three independent experiments, the MIC was the same across all experiments.

| Strain | MIC (μM) |
| --- | --- |
| BPTI-R A | 40 |
| BPTI-R B | 40 |
| BPTI-R C | 40 |
| Wildtype | 10 |
| ERG3 KO | 2.5 |
| GDA1 KO | 5 |
| NRP1 KO | 10 |
| PTK2 KO | 40 |
| SKY1 KO | 40 |

### Confirmation of mutation resistance

It was considered likely that the predicted missense or disruptive mutations in coding regions would have resulted in a loss of gene function. To test this hypothesis, single-gene knockouts of the mutated genes were retrieved from the yeast deletion set (Winzeler *et al.*, 1999) and antifungal assays were performed to assess whether gene deletion replicated the BPTI-resistant phenotype. Antifungal assays revealed that the single gene knockout mutants of *ptk2Δ* and *sky1Δ* were as resistant to BPTI as the BPTI-resistant isolates with an MIC of 40 μM. The knockout strains *gad1Δ* and *erg3Δ* were more sensitive to BPTI. The knockout *nrp1Δ* was inhibited at the same level as wildtype with an MIC of 10 μM (Table 4).

### BPTI and NaD1 act synergistically to inhibit BPTI and NaD1-resistant strains

BPTI and NaD1 have been reported to act synergistically in the inhibition of a range of pathogens (Bleackley *et al.*, 2017). Therefore, we assessed whether the NaD1-BPTI synergy still occurred if the strains were resistant to one of the peptides. The NaD1 and BPTI-resistant strains were tested against NaD1 and BPTI in a synergy assay and compared to wildtype (Figure 4). Wildtype had an average FIC synergy value of 0.23. The BPTI-resistant strains had an average FIC synergy value of 0.13 (Figure 4C), and the NaD1-resistant strains had an average FIC value of 0.19 (Figure 4F). Interestingly, the resistance of the BPTI resistant strains to BPTI was abolished upon addition of the lowest concentration of NaD1 (0.16 μM) (Figure 4A and C). Similarly, the resistance of the NaD1 strains to NaD1 was abolished in the presence of 0.16 μM BPTI (Figure 4D and F).

**Figure 4.**
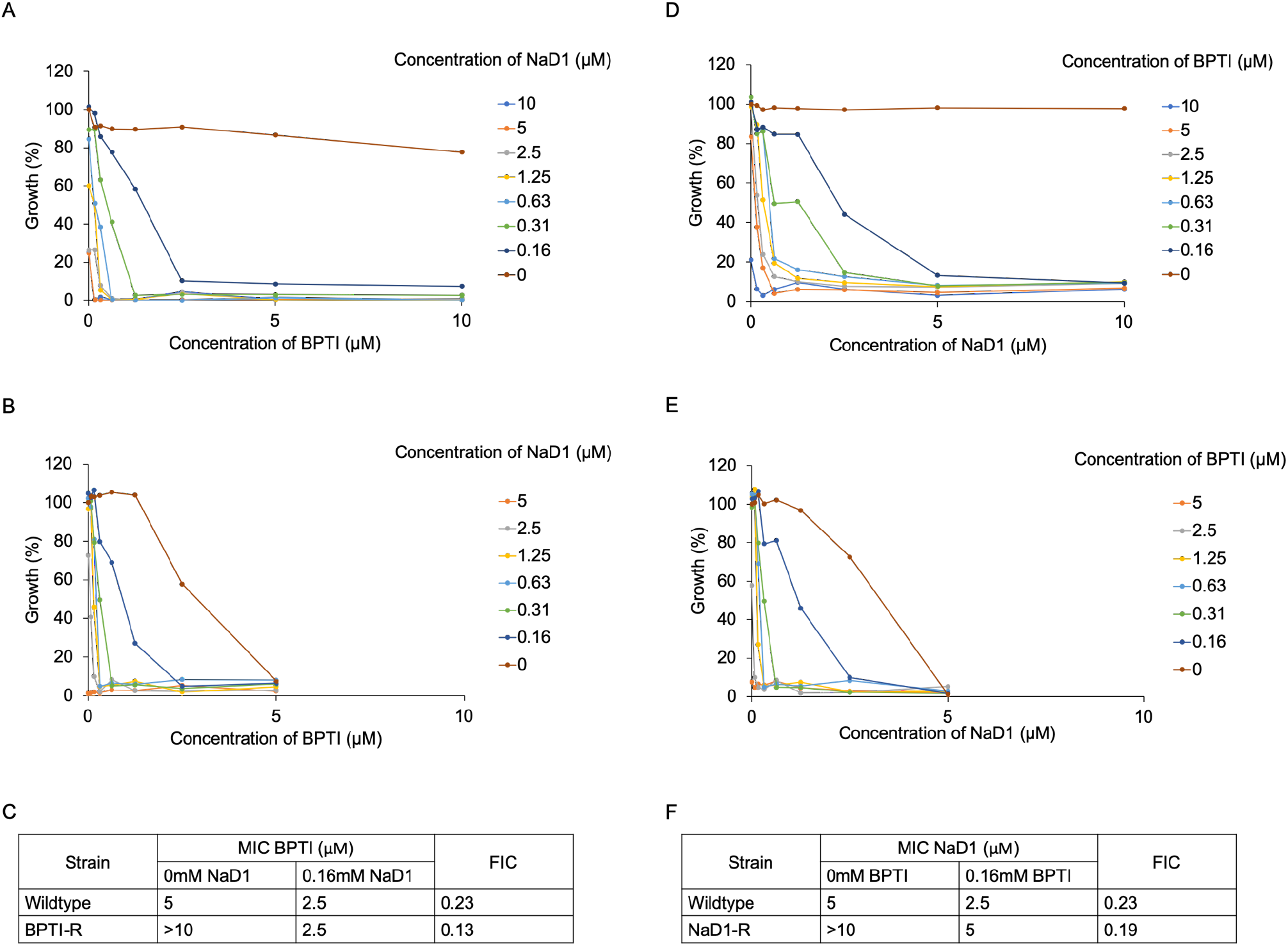
Synergy between NaD1 and BPTI on BPTI and NaD1 resistant strains is similar to wildtype. In the absence of NaD1, growth of BPTI-R was not inhibited by BPTI at concentrations up to 10 uM ((A) brown line) whereas growth of wildtype was fully inhibited at 5 uM BPTI ((B) brown line). However, when NaD1 was added at concentrations as low as 0.16 uM the growth inhibition of BPTI-R ((A) dark blue line) reverted to that observed for wildtype ((B) dark blue line). The MIC and FIC values for wildtype and BPTI-R in the presence and absence of the synergy partner are presented in a table (C). In the absence of BPTI, growth inhibition of NaD1-R was not inhibited by NaD1 at concentrations up to 10 uM ((D) brown line) whereas growth of wildtype was fully inhibited at 5 uM NaD1 ((E) brown line). When BPTI was added at concentrations as low as 0.16 uM the growth inhibition of NaD1-R ((D) dark blue line) reverted to that observed for wildtype ((E) dark blue line). The MIC and FIC values for wildtype and NaD1-R in the presence and absence of the synergy partner in are presented in a table (F). A comparison of the FIC, which is indicative of the strength of synergy, between wildytype, NaD1-R and BPTI-R (C and F) revealed that the level of synergy between NaD1 and BPTI is not affected by the mutations that lead to increased tolerance to each protein alone.

## Discussion

Aprotinin, also known as BPTI, was used during surgery to prevent blood loss in patients (Bidstrup *et al.*, 1989). Although BPTI is approved in some countries, it is restricted and the technological advances in medicine have produced better antifibrinolytic drugs (Van der Linden *et al*, 2001). We have been investigating another potential medical application for BPTI. BPTI has antifungal activity against the yeast S. *cerevisiae* with an MIC of 12.5 μg/mL, *C. neoformans* with an MIC of 9.4 μg/mL, and *C. albicans* with an MIC of 100 μg/mL (Table 1). BPTI did not have activity against the filamentous fungi *T. rubrum* and *A. fumigatus* at concentrations up to 200 μg/mL (Table 1). These results prompted the question of whether BPTI has the potential to be used as a treatment for systemic yeast infections. Such an application would depend on whether the yeast is able to develop resistance to BPTI quickly and how this would be managed. Here we showed that yeast do become resistant to BPTI after serial passaging in the presence of increasing amounts of peptide. Indeed resistance to BPTI developed more quickly than resistance to caspofungin and the plant defensin NaD1 that we described in an earlier publication (McColl *et al.*, 2018). Some BPTI resistant strains were cross resistant to other antifungal peptides, specifically LL37 and NaD2. However, the BPTI-resistant strains were still as susceptible to both NaD1, and caspofungin and they were more sensitive to Bac2a. That is, the resistance to BPTI was not due to a general improvement in fitness. The three strains that were resistant to BPTI were also more resistant to osmotic stress from elevated NaCl concentrations, and cell wall stress induced by calcofluor white. One strain was more resistant to hydrogen peroxide. Sequencing the genomes of the BPTI resistant strains and follow-up experiments using whole gene deletion strains revealed that mutations in *ptk2* and *sky1* are likely to be the primary source of resistance to BPTI.

BPTI exerts its antifungal activity by blocking Mg^2+^ uptake in to the cell and inhibiting growth (Bleackley *et al.*, 2014). Hexamine (III) cobalt chloride (HCC), a well characterized CorA Mg^2+^ transport inhibitor, inhibits cells at the same stages of the cell cycle as BPTI (Bleackley *et al.*, 2014) and results in a similar drop in cellular Mg ^2+^ levels. We thus assessed whether BPTI-resistant strains generated in this study were more tolerant to inhibition of magnesium uptake by assessing the effect HCC on their growth. The BPTI-resistant strains were 4-fold more tolerant to HCC with an MIC of 10 μM HCC compared to the wildtype strain which was fully inhibited at 2.5 μM. This 4-fold increase in resistance to HCC parallels the increase in resistance to BPTI observed in these strains. The similarities in the resistance to HCC indicate that resistance to BPTI is related to magnesium transport inhibition.

### Yeast develop resistance to BPTI more quickly than other antifungals

Resistance to BPTI developed quickly with 12 rounds of sub-culturing compared to 15 rounds for caspofungin and 21 rounds for resistance in NaD1 (Figure 1A). We attributed this relatively rapid development of resistance to two factors. The first is that BPTI is fungistatic, that is, it inhibits fungal growth but does not actively kill the fungus (Bleackley *et al.*, 2014) leaving a larger pool of living cells to develop mutations that confer resistance. Both NaD1 and caspofungin are fungicidal molecules. Fungicidal drugs are usually the preferred choice of treatment in the clinic because they act quickly and kill almost all cells (Kumar *et al*, 2018).The second reason is that BPTI inhibits fungal growth by blocking magnesium uptake by the membrane transporter Alr1p (Bleackley *et al.*, 2014). In contrast, NaD1 has a complex mechanism of action that involves: interaction with the fungal cell wall (van der Weerden *et al*, 2008), movement across the plasma membrane, induction of oxidative stress, and interaction with phosphatidylinositol 4,5 bisphosphate (Parisi *et al*, 2019). These processes lead to damage of the inner leaflet of the cell membrane and cell death within 10 min of exposure to NaD1 (Hayes *et al*, 2014; Payne *et al*, 2016; van der Weerden *et al*, 2010). Resistance to NaD1 develops more slowly compared to caspofungin because resistance to caspofungin can be achieved through point mutations in specific “hot spot” regions in the *fks1* gene, whereas resistance to NaD1 occurs through an accumulation of mutations in different genes (McColl *et al.*, 2018). It is likely that resistance to BPTI developed more quickly because there is only one component of the inhibitory mechanism, that is magnesium transport.

### BPTI resistant strains are also resistant to LL37 and NaD2 but not other antifungals

To assess how specific the resistance to BPTI was, the BPTI-resistant strains were tested against a range of other antifungals including; the antimicrobial plant defensin NaD1, another plant defensin from *Nicotiana alata* NaD2 (Dracatos *et al.*, 2014), the echinocandin caspofungin which is used in the clinic (McCormack & Perry, 2005), the human cathelicidin LL-37(Ordonez *et al*, 2014), and a linear variant of the bovine antimicrobial peptide bactenecin, Bac-2a (Hilpert *et al*, 2005) (Figure 2). There was no cross resistance associated with caspofungin, NaD1 or Bac2a. Therefore, resistance to BPTI would not provide cross protection against NaD1 or caspofungin as BPTI acts predominantly by blocking magnesium transport, which is a very different mechanism of action to NaD1 and caspofungin. However, the BPTI-resistant strains were more resistant to LL-37 as well as NaD2, but to a lesser extent. LL-37 associates with cell wall components of *C. albicans,* leading to cell membrane disruption (Burton & Steel, 2009; Tsai *et al*, 2014). We found that BPTI-resistant strains are also resistant to osmotic stress and calcofluor white, a cell wall stressor that binds to chitin (Figure 3A) which could explain the cross resistance to LL37. Similarly, identification of mutations in genes *sky1* and *ptk2*, which are responsible for the regulation of cation ion transport and homeostasis, would impact the ion gradients across the plasma membrane and therefore place stress on the membrane which in turn could influence susceptibility to LL-37. This pattern of cross resistance was not observed with the NaD1-resistant strains which were also resistant to other plant defensins, DmAMP1 and HXP4 (McColl *et al.*, 2018).

### Resistance to BPTI developed as the result of a deleterious mutation in one of two genes

Whole genome sequencing and SNV calling revealed a set of mutations that were present in the resistant strains, these were in the genes *sky1, ptk2, gad1, erg3,* and *nrp1.* It is likely that some of the mutations would result in a loss of gene function, particularly the mutations that were identified as deleterious. To test this hypothesis, single-gene knockouts corresponding to the mutated genes were retrieved from the yeast deletion set (Winzeler *et al.*, 1999) and antifungal assays were performed to assess whether gene deletion replicated the BPTI-resistant phenotype. Indeed, the single gene knockout mutants of of *ptk2Δ* and *sky1Δ* were as resistant to BPTI as the BPTI-resistant isolates at 40 μM. The BPTI-resistant strains A and C have a deleterious *ptk2* mutation (Gly469 to STOP) and BPTI-resistant strain B has a deleterious *sky1* mutation (Gln65 to STOP) (Table 1). The single gene knockout mutants of *gad1Δ* and *erg3Δ* were more sensitive to BPTI, this is likely because the mutations were either tolerated missense mutations or the mutations had an up-regulatory effect. The knockout *nrp1*Δ was inhibited the same as wildtype at 10 μM as expected because the mutations in nrp1p were unlikely to affect gene function. It was thus considered likely that all the resistance to BPTI was contributed by the mutations in *ptk2* or *sky1* because the *gad1*Δ, *erg3Δ* and *nrp1Δ* knockouts were not more resistant to BPTI. The mutations in *ptk2* and *sky1* both introduced an early stop codon, and thus would have the same effect as a gene knockout. Interestingly the BPTI-resistant strains also exhibited phenotypes that have been reported previously for *ptk2Δ* and *sky1Δ* strains. For example knockouts of *sky1* have been reported to be more resistant to hydrogen peroxide and calcofluor white (Brown *et al*, 2006), and we found that the BPTI-resistant strain B, with the *sky1* mutation, was more resistant to hydrogen peroxide and that this strain together with the other two BPTI-resistant strains were resistant to CFW (Figure 3A). Another screen of a yeast deletion library revealed that *sky1* and *ptk2* deletion mutants are resistant to sodium chloride, as we observed with the BPTI-resistant strains (Yoshikawa *et al*, 2009). Apart from the observed resistance to osmotic and cell wall stress, the BPTI-resistant strains had no other apparent fitness defects even in the presence of several abiotic stressors. This contrasts with the NaD1-resistant strains which had a decreased growth rate and were more sensitive to Calcofluor white and SDS (McColl *et al.*, 2018).

Bleackley and coworkers reported that BPTI interacts with the transporter Alr1p, blocking magnesium uptake into *S. cerevisiae* (Bleackley *et al.*, 2014). Surprisingly, we did not find any mutations in the *alr1* gene in the BPTI-resistant strains. *ALR1* is an essential gene and it is possible that mutations that would block BPTI binding are also deleterious to the essential function of the gene product in cellular Mg^2+^ uptake and therefore are not viable or have a serious fitness penalty. However, Ptk2p and Sky1p both function in cation homeostasis and transport, the mutations in these genes likely prevent the release of magnesium and other cations out of the cell, protecting the yeast from limiting intracellular magnesium levels that would be caused by BPTI blocking of Alr1p. Mutations in *sky1* and *ptk2* may be easier routes to increase magnesium levels, without a serious impact on cell vitality.

### BPTI and NaD1 retain synergy on resistant strains

Combination therapy with more than one antimicrobial is a potential mechanism to prevent the development of resistance in pathogenic microorganisms. In some cases, there is the added advantage that the two molecules work in synergy, reducing the inhibitory concentrations of the antimicrobial molecules below the levels predicted from their additive effect. We have reported previously that BPTI and NaD1 act synergistically to kill fungi (Bleackley *et al.*, 2017). Further investigation into the mechanism of synergy between NaD1 and BPTI revealed that the protease inhibitory activity of BPTI was not required for synergy and that BPTI also acted synergistically with other non-defensin antifungals (Bleackley *et al.*, 2017). It was hypothesized that protease inhibitors such as BPTI, influence stress response pathways that alter the ability of the fungus to respond to NaD1, and synergy could result from overloading of these stress response pathways when exposed to both NaD1 and BPTI. We therefore assessed whether the BPTI and NaD1-resistant strains were susceptible to the synergistic activity of NaD1 and BPTI. The strains that were resistant to NaD1 or BPTI had their sensitivity to the respective antifungal restored once a small amount of the opposing molecule was added (Figure 4). This is likely because the mutations associated with resistance are associated with mitigating the stress responses that occur after exposure to NaD1 or BPTI (Hayes *et al.*, 2014; McKenna, 2012). However, once the other peptide is added, these stress response pathways that normally enhance resistance are no longer effective. NaD1 resistant strains were sensitive to the cell wall stressor CFW (McColl *et al.*, 2018), whereas BPTI-resistant strains were resistant to CFW, this opposition in stress response may explain the rescue of antifungal synergy between BPTI and NaD1.

## Conclusion

Antimicrobial peptides are often touted as an attractive alternative to small molecule antimicrobials because they are more robust in terms of the potential for resistance to emerge (Mookherjee *et al*, 2020). Rapid development of resistance to the antifungal activity of BPTI would seem to contradict this idea but is likely to be a reflection of the superiority of fungicidal molecules as antifungals compared to fungistatic molecules. The fact that the resistance that was developed to either NaD1 or BPTI could be reverted by the addition of very low concentrations of the partner peptide in synergy assays indicates that if AMPs were developed for clinical use any increase in pathogen tolerance may be easy to combat through the use of combinatorial therapies that inactivate the stress response related resistance mechanisms.

## Supplementary figure list

**Supplementary figure A.**
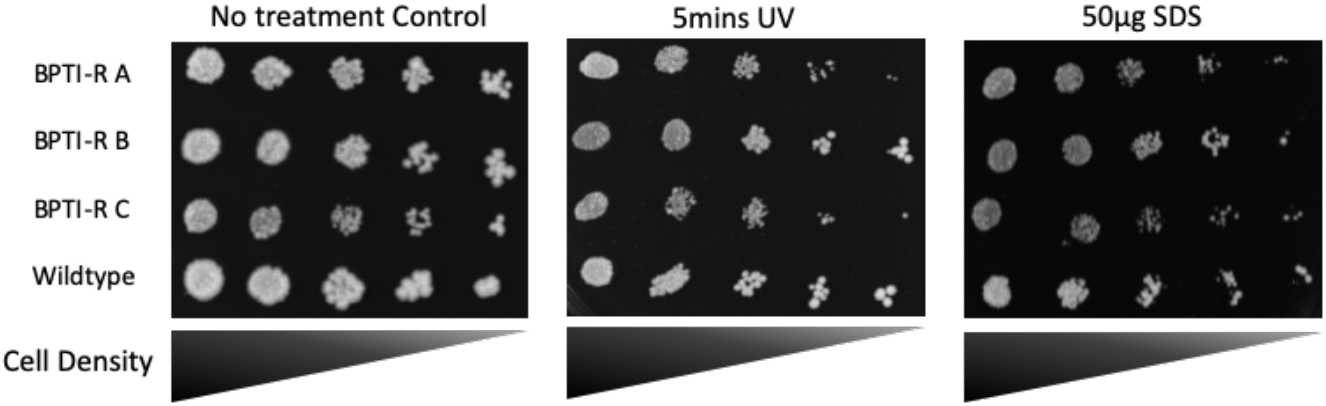
BPTI-resistant strains showed no difference in sensitivity to UV or SDS to the wildtype strain. (left) No treatment control plate that was incubated at the same time as the treatment plates. (middle) BPTI-resistant and wild-type *S. cerevisiae* BY4741 cells were diluted and spotted out on to YPD agar, before exposure to UV for 5min. There was no difference in sensitivity between the BPTI-resistant strains, control plate, and wild type, when exposed to UV. (right) There is no difference in sensitivity of the BPTI-resistant strains to SDS compared to the controls and wild type.

**Supplementary figure B.**
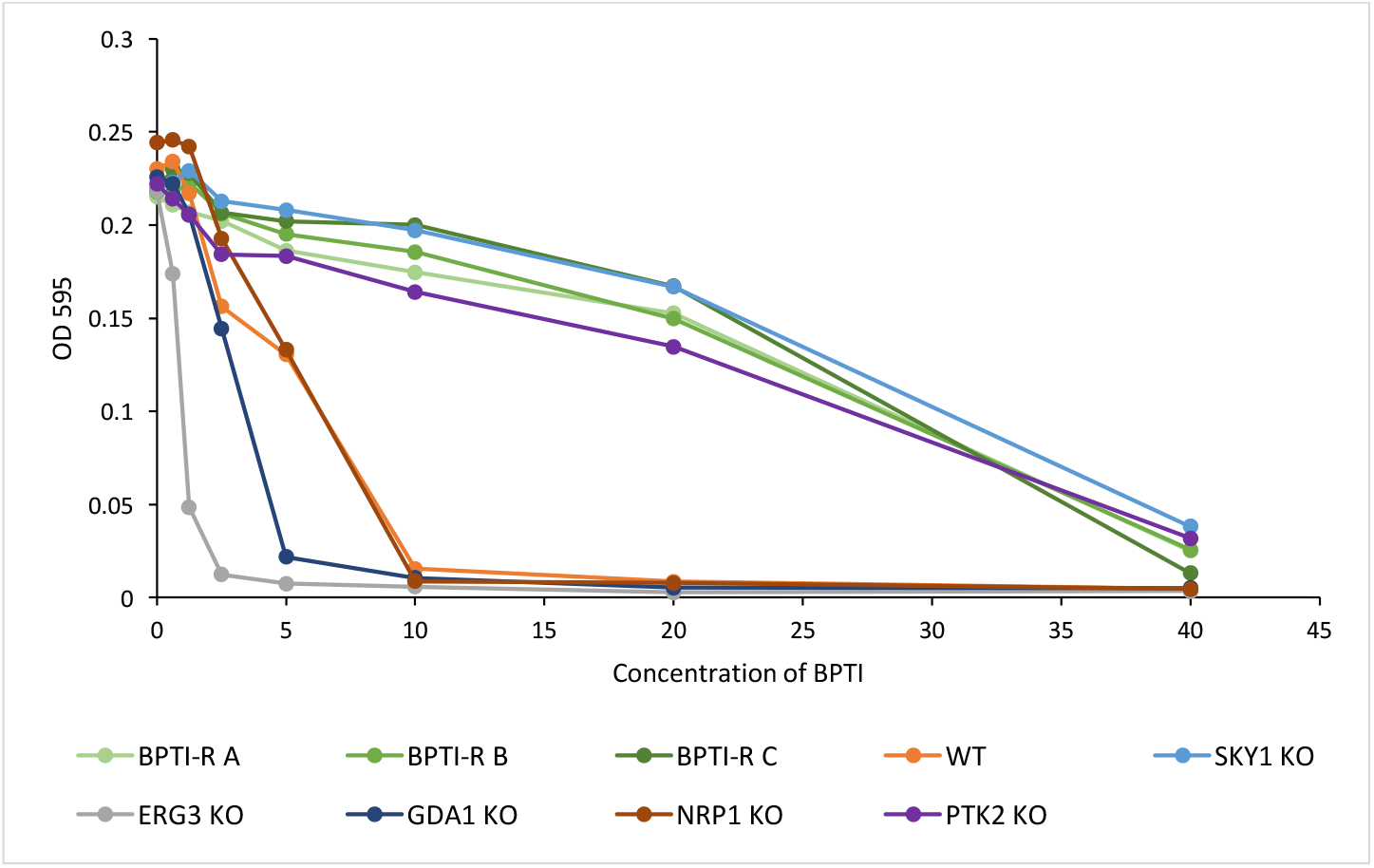
Comparison of BPTI activity against single-gene deletion strains representing key resistance variants. The activity of BPTI against BPTI-resistant strains A, B and C, wild-type *S. cerevisiae* BY4741 and the gene knock outs of ERG3, GDA1, NRP1, PTK2 and SKY1, the antifungal assay was performed in ½ PDB with the data being a representative example of three independent experiments, with all experiments having similar MIC.

